# 3D-Cell-Annotator: an open-source active surface tool for single cell segmentation in 3D microscopy images

**DOI:** 10.1101/677294

**Authors:** Ervin A. Tasnadi, Timea Toth, Maria Kovacs, Akos Diosdi, Francesco Pampaloni, Jozsef Molnar, Filippo Piccinini, Peter Horvath

## Abstract

**Summary:** Segmentation of single cells in microscopy images is one of the major challenges in computational biology. It is the first step of most bioimage analysis tasks, and essential to create training sets for more advanced deep learning approaches. Here, we propose 3D-Cell-Annotator to solve this task using 3D active surfaces together with shape descriptors as prior information in a fully- and semi-automated fashion. The software uses the convenient 3D interface of the widely used Medical Imaging Interaction Toolkit (MITK). Results on 3D biological structures (*e.g*. spheroids, organoids, embryos) show that the precision of the segmentation reaches the level of a human expert.

**Availability and implementation:** 3D-Cell-Annotator is implemented in CUDA/C++ as a patch for the segmentation module of MITK. The 3D-Cell-Annotator enabled MITK distribution can be downloaded at: www.3D-cell-annotator.org. It works under Windows 64-bit systems and recent Linux distributions even on a consumer level laptop with a CUDA-enabled video card using recent NVIDIA drivers.

## 1 Introduction

Multicellular 3D biological models, the so-called “-oids” (*e.g*. spheroids, organoids) are increasingly used as cellular models for drug screening and toxicology studies, since they represent physiological proxies of human tissues and can replace animal models (Zanoni *et al.*, 2016). Light-Sheet Fluorescence Microscopy (LSFM), confocal and multiphoton systems allow an in-depth observation of tissues in the size range of a few hundred microns. Despite the exponentially growing popularity of 3D models and systems to visualize - oids, few tools are available to analyse large aggregates at a single cell level (Carragher *et al*. 2018). In this work, we are focusing on the nuclei segmentation problem of multicellular aggregates as one of the most fundamental tasks of bioimage analysis, and the starting point of further phenotype-based statistics. Recently it has been shown that deep learning-based systems highly outperform classical image processing methods in 2D nuclei segmentation (Hollandi *et al*., 2019). However, these methods need accurate and large training sets, *i.e.* 3D annotated spheroids and embryos in our special case. Creating such ground truth datasets in 2D is mostly straightforward by drawing the contours of each cell on a 2D canvas. Similarly, the obvious extension of this approach to 3D would involve the annotation of each slice of the volume data. However, it is quite evident that this oversimplified method is not only time-consuming, but also leads to discontinuous object surfaces, while the quality of segmentation strongly depends on the chosen (orthogonal) plane. To overcome these problems, we designed 3D-Cell-Annotator. It provides an alternative for precise outlining of 3D shapes using a special type of active surface model (Molnar *et al*., 2017).

**Fig. 1:**
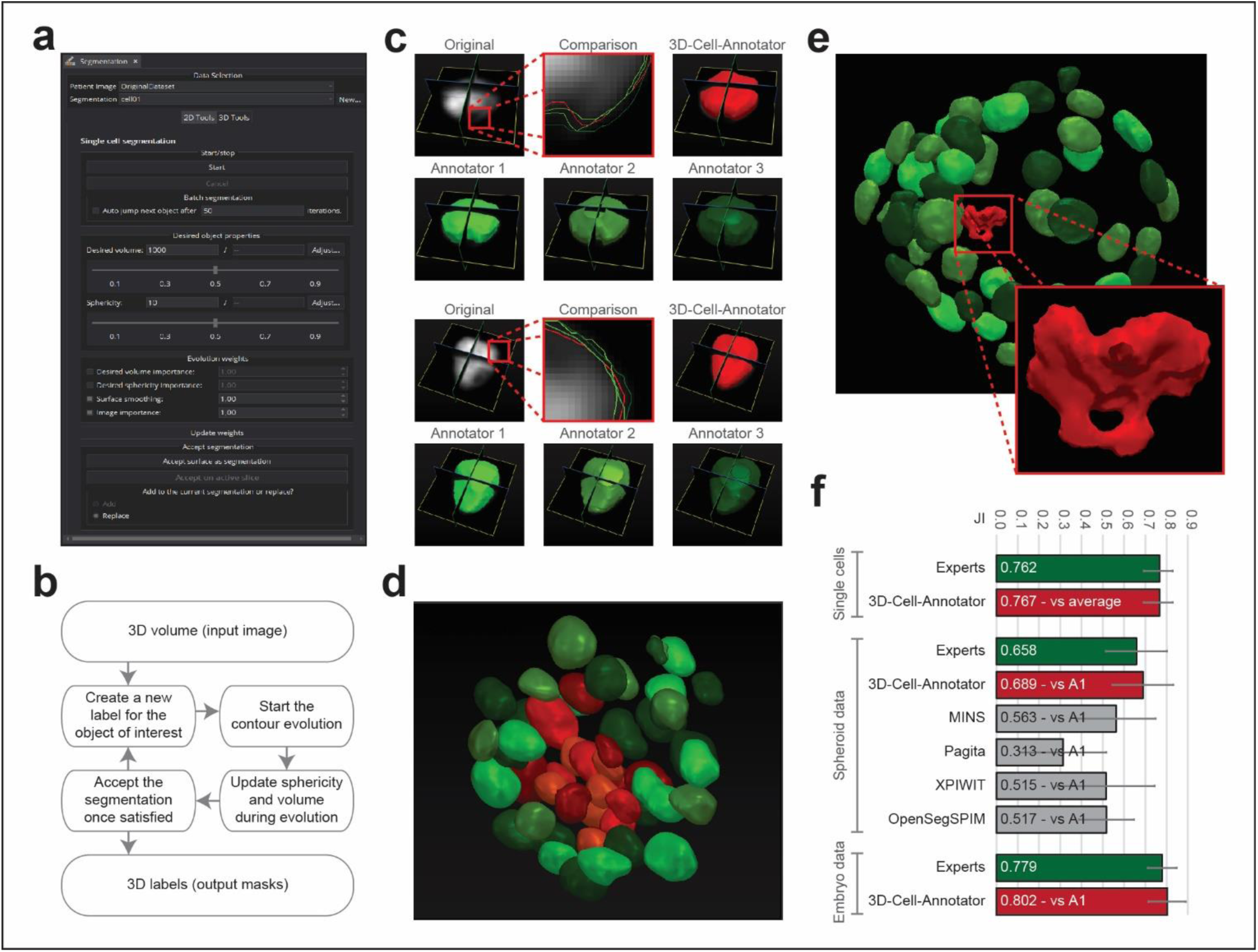
(**a**) 3D-Cell-Annotator Graphical User Interface (GUI). (**b**) Flow-chart of the segmentation approach. 3D-Cell-Annotator requires a 3D input image, typically a *z*-stack of sections acquired with a confocal/multiphoton/LSFM system. A label for each object is provided to start contour evolution. The user may adjust some parameters (*e.g*. volume, sphericity) in real time. Finally, the obtained segmentation can be exported as a 3D binary mask. (**c**) Manual segmentation dataset of single cells acquired by a confocal microscope, annotated by three experts. Despite the fact that the annotators are all experts in the field, the obtained segmentations slightly differ (green contours). However, those obtained by the proposed software does not vary significantly (red contour). (**d**) 3D segmentations of a cancer-derived multicellular spheroid, imaged with an LSFM at a single cell level. The segmentations were obtained by using 3D-Cell-Annotator. For visual purposes, we distinguish the cells in the inner core that in large aggregates are typically senescent/necrotic cells (shown in red), from the cells in the outer shell that are the highly proliferative ones (shown in green). (**e**) 3D-Cell-Annotator can be used to extract cells with a special phenotype, such as mitosis or apoptosis, as shown in the magnified cell (red). (**f**) Jaccard Indices (JI), comparing the 3D masks obtained manually by the human expert annotators, by four other freely available tools and by the proposed algorithm on the datasets shown in (c), (d) and (e). 3D-Cell-Annotator outperformed every other software and reached an expert-like precision. A1 stands for *Annotator 1*; JI for *Jaccard Index*.

## 2 The proposed software

Instead of the classical slice-by-slice manual annotation approach, we propose a real 3D active surface-based solution with shape priors called *3D selective segmentation* to obtain accurate single-cell annotation (Molnar *et al*., 2017). Because of the pure 3D nature of our method, the spatial dependencies across all dimensions are considered by the algorithm, thus most of the time-consuming hand-drawing work may be eliminated. 3D-Cell-Annotator is distributed as a module of the widely used Medical Imaging Interaction Toolkit (MITK, Nolden *et al.*, 2013). Active surface models are computationally complex and expensive, therefore our model was targeted to Graphic Processing Unit (GPU), implemented in the NVidia CUDA framework to provide a speed increase of several orders of magnitude compared to classical CPU implementations. Annotation can be provided cell-by-cell manually or automatical y by placing initial seedpoints. While the general active surface algorithm may output clusters of objects when multiple cells share boundaries, the proposed selective active surface applies forces to fulfil shape descriptor values provided by the user (Molnar *et al*., 2017). Two such descriptors are used: sphericity and the volume of the object. Mathematical foundations are explained in **Supplementary Material 1**. Briefly, sphericity is a measure of the degree to which an object’s shape is similar to a sphere. The volume prior is used to stop the surface to over-/undergrow. These prior parameters can be fine-adjusted with high precision during surface evolution to obtain segmentation at a single cell level. A fully automated batch segmentation mode is also available.

## 3 Results

To evaluate 3D-Cell-Annotator we used (*a*) a confocal dataset of 77 *z*-stacks, each containing one single cell (Poulet *et al*. 2014); (*b*) an LSFM dataset used in Gole *et al*. 2016, representing a low intensity contrast multicellular spheroid composed of 52 cells; and (*c*) a mouse embryo dataset containing 56 cells acquired by a confocal microscope presented in Saiz *et al*. 2016. All the datasets we used in this work are publicly available (**Supplementary Material 2**). We computed the Jaccard Index (JI) for the segmentations obtained by 3D-Cell-Annotator compared to other tools, as well as to manual segmentations executed by expert annotators. Our binary masks were diluted by a 1 pixel radius sphere to overcome the problem of active surface discretisation. The MATLAB (The MathWorks, Inc., MA, USA) code to compute the JI for 3D binary masks for multiple objects is provided as **Supplementary Material 3**.

### 3.1 Single-cell dataset

To evaluate the segmentation quality of 3D-Cell-Annotator we used a dataset of 77 *z*-stacks, each containing a single cell. The masks obtained by 3D-Cell-Annotator were compared with manual “ground truth” segmentation. In order to obtain the ground truth for this dataset, we asked three expert annotators to segment the 77 *z*-stacks. For each stack we averaged their 3D segmentations by utilizing ReViMS (Piccinini *et al*. 2017, De Santis *et al*. 2019), a 3D extension of STAPLE (Warfield *et al*. 2004). We segmented the cells in a semi-automatic way by manually initializing with three contours on the three orthogonal axes, and then visually adjusting the parameters during surface evolution. The time spent by each human annotator on manually segmenting the 77 *z*-stacks was approximately 8 hours, whilst the time needed to obtain the same segmentations with 3D-Cell-Annotator was around 4 hours. The average JI between the expert annotators was 0.76. The JI between the annotators’ average segmentations and 3D-Cell-Annotator was 0.77. The accuracy of 3D-Cell-Annotator reaches the level of an expert human annotator. The JI values calculated for each *z*-stack are reported in **Supplementary Material 4**.

### 3.2 Spheroid and embryo datasets

To demonstrate the usability and accuracy of 3D-Cell-Annotator for the analysis of real multicellular aggregates, we used a spheroid dataset imaged with an LSFM. The spheroid is composed of 52 cells (Gole *et al*. 2016). We compared the segmentations presented by two expert annotators with those obtained using 3D-Cell-Annotator and other four freely available tools: MINS (Lou *et al*. 2014, Saiz *et al*. 2016), Pagita (Gul-Mohammed *et al*. 2014), XPIWIT (Bartschat *et al*. 2015), and OpenSegSPIM (Gole *et al*. 2016). A brief description of these tools is available in **Supplementary Material 5**. The average JI for the comparison of the performance of human experts was 0.66; the average JI comparing 3D-Cell-Annotator to the human annotators separately was 0.68 and 0.69, respectively, whilst none of the other tools performed better than 0.57 (**Supplementary Material 6**). T us, it can be concluded that 3D-Cell-Annotator outperforms the analysed tools and its accuracy reaches the level of human experts’. In order to confirm this latter claim, we tested the software on another publicly available dataset, namely on a dataset of a mouse embryo consisting of 56 cells. Two expert operators segmented the cells manually in approximately 6 hours, whilst the time needed to obtain the same segmentations with 3D-Cell-Annotator was 3 hours. The average JI between human experts was 0.78; be ween 3D-Cell-Annotator and both the human annotators was 0.80 (**Supplementary Material 7**). Again, these results confirm that 3D-Cell-Annotator offers an accuracy level reaching that of a human expert.

## 4 Conclusions

3D-Cell-Annotator provides a user friendly and precise solution for segmenting single cells in 3D cell cultures imaged with confocal microscope or LSFM, even for large datasets with touching cells and suboptimal imaging conditions, like in the case of spheroids, organoids and embryos. Reaching the accuracy of an expert human annotator, and permitting to save at least half of the time spent, 3D-Cell-Annotator is an optimal solution for generating trining tests for more advanced machine learning approaches. Further improvements will include the implementation of different approaches for cell splitting. User manual, video tutorials, and all the masks discussed in this work are available at: www.3D-cell-annotator.org.

## Acknowledgements & Funding

The authors would like to thank Laurent Gole (Institute of Molecular and Cell Biology, A*STAR, Singapore), Anna-Katerina Hadjantonakis and Nestor Saiz (Sloan Kettering Institute, New York, USA) for material and important information shared about tools for 3D cell segmentation; Máté Görbe (BRC, Hungary) for creating the website and videos; Gabriella Tick (BRC, Hungary) for recording the video tutorial narration; Dora Bokor, PharmD (BRC, Hungary) for proofreading the manuscript. ET, TT, MK, AD, JM, FP, and HP acknowledge support from the LENDULET-BIOMAG Grant (2018-342) and from the European Reional Development Funds (GINOP-2.3.2-15-2016-00026, GINOP-2.3.2-15-2016-00037); FP acknowledges support from the Union for International Cancer Control (UICC) for a granted UICC Technical Fellowship (ref: UICC-TF/19/640197).

