## Supplementary Material for "3D-Cell-Annotator: an open-source active surface tool for single cell segmentation in 3D microscopy images"

**Supplementary Material 1**

### 3D-Cell-Annotator technical details

#### 1 Introduction

In this supplementary, we introduce the theoretical foundations for the methods used in the software. 3D-Cell-Annotator is based on an extended version of active contours called selective active contour model.[1] The selective active model introduces two different priors, a volume and a shape prior, these priors are proposed in the 3D-Cell-Annotator to extract single cells from clusters. Two different data terms are tested, the first one is the simplest anisotropic edge detector while the other one considers the environment of the contour to better find the boundary of the object, called the local region data term. The implementation is done in a level set framework. Since one of the most fundamental limitation of the level set method is its numerical instability, we proposed a new method called balanced phase field model, that regularizes the level set in a narrow band in every iteration step to achieve numerical stability during the segmentation.[2] We briefly discuss the selective and the balanced phase field models here since these are the main building blocks of the algorithm used by the 3D-Cell-Annotator.

#### 2 Selective active contours in 3D

The 2D selective active contour model was introduced to segment objects by their size and shape.[1, 3]. We chose the level set framework as the implementation of the model as it proved to be useful for implementing interfacial problems like active contour models. [4]

##### 2.1 Notations

Surfaces are denoted by $\mathbf{S} \subseteq \mathbb{R}^3$ or $\mathbf{S}(u,v) \in \mathbb{R}^3$ in parameterized form, where $u$ and $v$ are surface parameters. $\mathbf{S}_u(u,v), \mathbf{S}_v(u,v) \in \mathbb{R}^3$ are partial derivatives wrt surface parameters (u,v), providing local (covariant) basis for the vectors of the tangent plane at $\mathbf{S}(u,v)$. Recall that $\mathbf{S}_u \times \mathbf{S}_v$ is normal to the surface. Assuming $\mathbf{S_u}, \mathbf{S}_v\mathbf{n}$ constitute a right handed basis, the inward pointing unit normal $\frac{\mathbf{S}_u \times \mathbf{S}_v}{|\mathbf{S}_u \times \mathbf{S}_v|}$ is denoted by $\mathbf{n}$. The sum curvature of the surface is denoted by $K$, while $K_G$ is the Gaussian curvature. The integral $\int dS = \int \sqrt{|\mathbf{S}_u|^2|\mathbf{S}_v|^2 - (\mathbf{S}_u \cdot \mathbf{S}_v)^2} dudv$ gives the surface area and $\int dV = -\frac{1}{6}\int \mathbf{S} \cdot (\mathbf{S}_u \times \mathbf{S}_v) dudv$ gives the volume of a surface $\mathbf{S}$, where $dS$ and $dV$ are the surface and volume element respectively.

Level set functions are denoted by $\phi = \phi(t, \mathbf{x})$, where $t \in \mathbb{R}$ and $\mathbf{x} = (x_1, x_2, x_3) \in \mathbb{R}^3$ are the time and space variables respectively. According to this, $\phi_t$ denotes the partial derivative with respect to the time and $\nabla\phi$ denotes the spatial gradient $\nabla\phi = (\phi_{x_1}, \phi_{x_2}, \phi_{x_3})$. The Hessian matrix of $\phi$ is denoted by $\mathbf{H}(\phi) = (\phi_{x_i x_j})_{1 \leq i,j \leq 3}$.

#### 2.2 The selective functional

There are four terms included in the functional of the selective active contour model. Below we provide a concise summary. For the details, see the original paper. [1]

##### 2.2.1 Volume prior

The volume prior prefers objects having certain size and it is denoted by $V_0$.

$$\mathcal{V}(\mathbf{S}) = \frac{1}{kV_0^k}\left(\int dV - V_0\right)^k \tag{1}$$

The $k$ is an arbitrary integer. If it is set up to 2 then the contour try to have a volume exactly $V_0$ while if it is set up to 3, it has an inflection at $V_0$ therefore it prefers a volume of 0 except at $V_0$ where the term has no effect.

##### 2.2.2 Shape prior

The shape prior penalizes the deviation of the current surface from the preferred shape $p$ (plasma or amoeba value). The currently implemented shape prior is called *sphericity* in the 3D-Cell-Annotator but essentially it is the surface/volume ratio of the surface that is $p = \frac{area^{\frac{3}{2}}}{volume}$. The plasma value is minimal for the sphere, that is exactly $p = 3\sqrt{4\pi} \approx 10.6$. We want the sphericity to have a value of 1.0 for the sphere so the sphericity is given by $p - 10.6$.

The shape prior that considers the surface volume ratio is then:

$$\mathcal{S}(\mathbf{S}) = \frac{1}{2V_0^2}\left[\left(\int dS\right)^{\frac{3}{2}} - p\int dV\right]^2 \tag{2}$$

##### 2.2.3 Smoothness term

A curvature based smoothness term is applied to prevent the instability of the surface called Euler elastica. For the details, consult the original paper. [1]

$$\mathcal{E}(\mathbf{S}) = \frac{1}{2}\int K^2 dS, \tag{3}$$

where the $K$ is the sum curvature of the surface at a given point.

##### 2.2.4 Data term

Two different data term is tested, but it can be chosen from a wide range of possible ones. The first is an edge detector:

$$\mathcal{D}_{\mathcal{E}}(\mathbf{S}) = \int \nabla I \cdot \mathbf{n} dS, \tag{4}$$

where the $I$ is the image.

The second one is a region based data term. It considers the mean intensity in rectangular prism shaped local region positioned both in the inner and the outer part of the contour. In our setting the functional maximizes the intensity difference between the inner and outer part. If this one is used, then the algorithm has three more parameters defining the size of the prism (width, height and depth).

$$\Phi(\mathbf{S}, \mathbf{n}) = \frac{1}{4pqr} \left( \int_{\Re^+} I(\mathbf{p}) dV - \int_{\Re^-} I(\mathbf{p}) dV \right), \tag{5}$$

where $dV = d\xi d\zeta d\eta$, $\xi \in [-p, p]$, $\zeta \in [-q, q]$ and $\eta \in [0, r]$. $\mathbf{p}$ is in the local coordinate system, therefore $\mathbf{p} = \mathbf{S} + \xi \mathbf{e_1} + \zeta \mathbf{e_2} + \eta \mathbf{n}$ as it is visualised in fig. (1).

Therefore, we can use the local region as a data term in the selective model:

$$\mathcal{D}_{\mathcal{R}}(\mathbf{S}) = \int \Phi(\mathbf{S}, \mathbf{n}) dS. \tag{6}$$

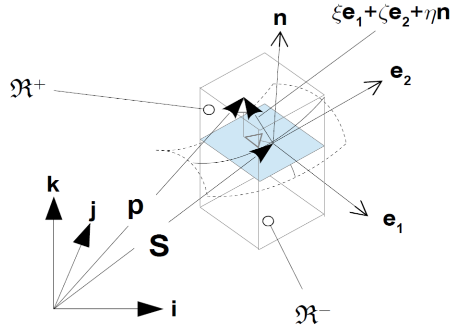


Figure 1: Visualisation of the local region in 3D. The local Cartesian coordinate system of the region is centered at the surface point $\mathbf{S}$ while the orientation is determined by the unit normal vector of the surface $\mathbf{n}$ and the unit basis vectors $\mathbf{e_1}, \mathbf{e_2}$ of the tangential plane of the surface.

##### 2.2.5 Putting all together: the composite functional

The composite functional therefore consists of the previously introduced terms therefore it becomes to:

$$\mathcal{L} = \alpha\mathcal{D} + \beta\mathcal{S} + \gamma\mathcal{V} + \delta\mathcal{E}, \tag{7}$$

where each term has an arbitrary real weight that can be controlled from the interface of the 3D-Cell-Annotator. The $\mathcal{D}$ can be $\mathcal{D}_{\mathcal{E}}$ or $\mathcal{D}_{\mathcal{R}}$ either depending on which data term is used.

#### 2.3 The Euler-Lagrange equation for the functional

The extremal surface of the functional above can be found by solving the corresponding Euler-Lagrange equations. In our case (3D surfaces in the level set framework) they have the form:

$$|\mathbf{S}_u \times \mathbf{S}_v| Q\mathbf{n} = \mathbf{0}, \tag{8}$$

where $Q$ is a scalar field with the functional derivatives: $Q = \alpha Q_{\mathcal{D}_{\mathcal{E}}} + \beta Q_{\mathcal{S}} + \gamma Q_{\mathcal{V}} + \delta Q_{\mathcal{E}}$.

That is, for the volume prior, we have $Q_{\mathcal{V}} = -\frac{1}{V_0^3}(\int dV - V_0)^2$ ($V_0$ is the target volume), for the shape prior (that takes sphericity into consideration) $Q_{\mathcal{S}} = \left[\left(\int dS\right)^{\frac{3}{2}} - p\int dV\right]\left[p - \frac{3}{2}K\left(\int dS\right)^{\frac{1}{2}}\right]$, $p$ is the (unnormalized) target plasma value ($p = \frac{surface^{\frac{3}{2}}}{volume}$), for the data term, we have $Q_{\mathcal{D}_{\mathcal{E}}} = \Delta I$ and for the smoothness term we have $Q_{\mathcal{E}} = \frac{1}{2}K^3 - 2K_G K + \nabla \cdot \nabla K$, where $K_G$ is the Gaussian curvature. The Euler-Lagrange of $\mathcal{D}_{\mathcal{R}}$ is slightly more complicated.

### 3 Level set regularization: the Balanced Phase Field model

#### 3.1 Notations

In the level set framework, the representation of contours is given by a level set function of two variables $\phi(x, y)$. The quantities of the segmentation problem are extracted from this function, such as the unit normal vector $\mathbf{n} = \frac{\nabla\phi}{|\nabla\phi|}$ or the curvature $\kappa = -\nabla \cdot \left(\frac{\nabla\phi}{|\nabla\phi|}\right)$ where $\nabla$ is the gradient operator and "$\cdot$" stands for the scalar (dot) product, *i.e.* $\nabla \cdot \mathbf{v}$ is the divergence of the vector field $\mathbf{v}$.

#### 3.2 From the phase field model to the balanced model

Since one of the fundamental problems of the level set method is its numerical instability, one should care about the numerical errors by using some regularization method explicitly by not letting the active contour model to deform the level set from the signed distance function too much. We experienced severe issues with our selective model when we computed the curve evolution in level set framework, and the tested approximate solutions did not solve the issue, while the more accurate methods made the algorithm too slow to be practical in reality. A simple observation is that it is not needed to maintain the whole level set during the evolution since we always consider the derivatives in the neighborhood of the interface. Therefore it is enough to maintain the stability in this neighborhood called narrow band. Therefore we chose to apply a modified phase field solution to regularize the level set in every iteration. Using such a model, we treat the level set as a phase field, where the inner part of the contour is encoded with a value 1 while the outer is with $-1$. Between the two, there is a *phase transition* with a necessary zero-crossing that models the contour. The goal is to maintain a smooth phase transition that is similar to the signed distance property of the level set near the contour. Furthermore, using the modified phase field model called balanced phase field, the authors does not only propose a solution to the numerical issues of the level sets but also eliminates the effect of the regularization method on the active model. In the following, we discuss the most important aspects of the model for the two dimensional case starting from the original phase field model and introducing the ideas behind the improved balanced phase field. For the details, and the intermediate steps of the derivation, consult [2].

In the original functional we had:

$$\int\int_{\Omega} \frac{D_0}{2}|\nabla\phi|^2 + \lambda_0\left(\frac{\phi^4}{4} - \frac{\phi^2}{2}\right)dA. \tag{9}$$

The solution of (9) is a scalar field $\phi$ with values $\pm 1$ and a phase transition between the two values representing the narrow band of the contour. It is possible to embed the functional (9) to the active contour model directly by extending its functional with it, however it may lead to an extremely complex system considering its analysis. Instead of applying the phase field directly, one can use it in a shape maintenance role: before the next evolution step, the Euler-Lagrange equation associated with (9) can be solved independently, providing a regularized narrow band to the next iteration step.

However, a careful analysis shows that applying this functional to the level sets, it has a serious side effect on the active model by producing a curvature driven motion. In order to fix this issue, a Laplacian smoothness term is introduced and the original functional becomes the following:

$$\int\int_{\Omega} \frac{D}{2}(\Delta\phi)^2 + \lambda\left(\frac{\phi^4}{4} - \frac{\phi^2}{2} + \frac{1}{4}\right)dA. \tag{10}$$

However the proposed functional (10) still have a curvature dependent term and therefore produces the same effect when applied. It was shown in the paper, that using the combination of the smoothness terms from the previous functionals in a new one, under special conditions, the curvature dependency can be almost fully eliminated. The new functional then becomes the following:

$$\int\int_{\Omega} \frac{D}{2}|\Delta\phi|^2 - \frac{D_0}{2}|\nabla\phi|^2 + \lambda\left(\frac{\phi^4}{4} - \frac{\phi^2}{2} + \frac{1}{4}\right)dA. \tag{11}$$

The conditions needed to satisfy in order to cancel the effect of the curvature in the functional (11) depend on the width of the phase transition ($w$):

$$\lambda w^4 - 24D_0 w^2 - 720D = 0, \tag{12}$$

and

$$-D_0\frac{3}{w} + D\frac{48}{w^3} = 0. \tag{13}$$

In order to satisfy the conditions (12) and (13), one should compute the parameters of (11) as a function of the wished width of the phase transition. Therefore we first should determine the width of the phase transition needed. This depends on the highest order of derivatives ($n$) used in the active model since we have to consider at least $n+1$ points around the contour if we use the central difference schemes. For safety reasons, $w$ should be at least $2n+1$ to make sure that the $n$ neighborhood around the contour approximates the signed distance function enough. If we decided about the $w$ parameter, we can set up $D_0$ arbitrary, in this case to 1 for simplicity. Then, solving the conditions above, for the remaining parameters we get:

$$D_0 = 1, D = \frac{w^2}{16}, \lambda = \frac{21}{w^2}. \tag{14}$$

By summing up, the associated Euler-Lagrange functional with the parameters only depending on the width of the phase transition is:

$$\frac{w^2}{16}\Delta\Delta\phi + \Delta\phi + \frac{21}{w^2}\left(\phi^3 - \phi\right) = 0. \tag{15}$$

The (15) can be implemented as simply as applying a (4-th order linear) filter modified by a point-wise value of the nonlinear (cubic) term for the grid points of the phase field lattice.

### 4 Implementation details

We briefly discuss the implementation of the selective algorithm and the 3D-Cell-Annotator software.

#### 4.1 The selective model with the balanced phase field reinitialization model

We solve the Euler-Lagrange equations in the level set framework, therefore the following quantities can be substituted:

$$\mathbf{S}_u \times \mathbf{S}_v \mapsto \nabla\phi, \quad \mathbf{n} \mapsto \frac{\nabla\phi}{|\nabla\phi|}, \tag{16}$$

while the curvatures are computed as:

$$K \mapsto -\nabla \cdot \frac{\nabla\phi}{|\nabla\phi|}, \quad K_G \mapsto |\nabla\phi|^{-4} \begin{vmatrix} \mathbf{H}(\phi) & \nabla\phi^T \\ \nabla\phi & 0 \end{vmatrix}. \tag{17}$$

For minimizing the functionals a simple gradient descent is used. The finite differences are used to compute the derivatives numerically.

##### 4.2 The software

The selective model is targeted to the CUDA architecture in C++. The level set values are only computed where the surface is located in the current iteration. The algorithm is distributed as a shared library and a new Tool is created in the MITK that implements the communication between the shared library and the MITK.

**Supplementary Material 2**

All the datasets used in this work are publicly available and are described in other scientific papers:

1. The confocal dataset consists of 77 *z*-stacks containing a single-cell in each image stack. It was described by Poulet *et al*. 2014 and is publicly available at:
   *https://www.gred-clermont.fr/media/WorkDirectory.zip*

2. The LSFM dataset represents a small multicellular spheroid of 52 cells, used in Gole et al. 2016. It is publicly available under the name of "Neurosphere_Dataset" at:
   *http://opensegspim.weebly.com/download.html*

3. The confocal mouse embryo dataset contains 56 cells. It was used by Saiz et al. 2016. It is named "4May15FGFRionCD1_SU54_LM1.lsm" and it is publicly available in the native microscopy format (*i.e.* Zeiss ".lsm" format) at:
   *https://figshare.com/articles/Raw_images_for_Saiz_et_al_2016/3767976/1*
   In our analysis, we just considered the blue channel of this dataset, that is the one related to nuclear staining.

#### Supplementary Material 3

The MATLAB (The MathWorks, Inc., MA, USA) code to compute the JI comparing 3D binary masks containing multiple objects, is provided together with some Utils at:

*http://www.3d-cell-annotator.org/download.html*

#### Supplementary Material 4

**Table 2.** Confocal dataset of 77 single-cell *z*-stacks: Jaccard Index (JI) values

| | **Annotator 1 vs Annotator 2** | **Annotator 1 vs Annotator 3** | **Annotator 2 vs Annotator 3** | **Annotators average vs 3D-Cell-Annotator (semi-automatic modality)** |
|---|---|---|---|---|
| SINGLE CELL JI | | | | |
| | 0.875 | 0.850 | 0.912 | 0.880 |
| | 0.873 | 0.844 | 0.894 | 0.868 |
| | 0.871 | 0.839 | 0.887 | 0.850 |
| | 0.868 | 0.838 | 0.876 | 0.848 |
| | 0.864 | 0.837 | 0.875 | 0.846 |
| | 0.864 | 0.833 | 0.870 | 0.846 |
| | 0.863 | 0.828 | 0.870 | 0.845 |
| | 0.859 | 0.822 | 0.868 | 0.844 |
| | 0.854 | 0.817 | 0.857 | 0.842 |
| | 0.853 | 0.808 | 0.854 | 0.840 |
| | 0.851 | 0.806 | 0.854 | 0.835 |
| | 0.851 | 0.804 | 0.851 | 0.832 |
| | 0.850 | 0.803 | 0.849 | 0.830 |
| | 0.849 | 0.801 | 0.847 | 0.830 |
| | 0.848 | 0.798 | 0.839 | 0.829 |
| | 0.848 | 0.785 | 0.836 | 0.828 |
| | 0.844 | 0.785 | 0.831 | 0.825 |
| | 0.844 | 0.782 | 0.831 | 0.823 |

| | | | |
|---|---|---|---|
| 0.844 | 0.779 | 0.830 | 0.822 |
| 0.842 | 0.776 | 0.828 | 0.820 |
| 0.839 | 0.768 | 0.822 | 0.820 |
| 0.838 | 0.759 | 0.822 | 0.816 |
| 0.836 | 0.757 | 0.820 | 0.812 |
| 0.834 | 0.755 | 0.818 | 0.810 |
| 0.833 | 0.754 | 0.817 | 0.808 |
| 0.832 | 0.750 | 0.816 | 0.801 |
| 0.831 | 0.743 | 0.814 | 0.799 |
| 0.828 | 0.743 | 0.809 | 0.796 |
| 0.824 | 0.739 | 0.808 | 0.796 |
| 0.824 | 0.738 | 0.805 | 0.795 |
| 0.823 | 0.738 | 0.803 | 0.794 |
| 0.820 | 0.737 | 0.801 | 0.791 |
| 0.819 | 0.736 | 0.799 | 0.789 |
| 0.818 | 0.735 | 0.798 | 0.785 |
| 0.815 | 0.732 | 0.796 | 0.785 |
| 0.814 | 0.727 | 0.796 | 0.782 |
| 0.814 | 0.726 | 0.795 | 0.782 |
| 0.809 | 0.723 | 0.789 | 0.780 |
| 0.808 | 0.722 | 0.788 | 0.780 |
| 0.805 | 0.719 | 0.788 | 0.778 |

| | | | |
|---|---|---|---|
| 0.804 | 0.716 | 0.784 | 0.777 |
| 0.799 | 0.701 | 0.781 | 0.772 |
| 0.794 | 0.700 | 0.776 | 0.766 |
| 0.793 | 0.698 | 0.775 | 0.765 |
| 0.787 | 0.694 | 0.775 | 0.762 |
| 0.785 | 0.693 | 0.772 | 0.761 |
| 0.784 | 0.689 | 0.767 | 0.747 |
| 0.783 | 0.685 | 0.767 | 0.746 |
| 0.781 | 0.681 | 0.765 | 0.743 |
| 0.778 | 0.681 | 0.765 | 0.741 |
| 0.776 | 0.679 | 0.764 | 0.738 |
| 0.776 | 0.675 | 0.761 | 0.737 |
| 0.768 | 0.671 | 0.760 | 0.737 |
| 0.768 | 0.669 | 0.757 | 0.735 |
| 0.760 | 0.664 | 0.757 | 0.733 |
| 0.757 | 0.663 | 0.753 | 0.730 |
| 0.751 | 0.653 | 0.745 | 0.724 |
| 0.744 | 0.652 | 0.744 | 0.723 |
| 0.744 | 0.652 | 0.742 | 0.717 |
| 0.743 | 0.645 | 0.737 | 0.712 |
| 0.742 | 0.643 | 0.735 | 0.710 |
| 0.742 | 0.636 | 0.735 | 0.709 |

| | | | | |
|---|---|---|---|---|
| | 0.739 | 0.635 | 0.726 | 0.707 |
| | 0.736 | 0.631 | 0.723 | 0.704 |
| | 0.733 | 0.630 | 0.712 | 0.698 |
| | 0.726 | 0.626 | 0.712 | 0.697 |
| | 0.724 | 0.613 | 0.709 | 0.696 |
| | 0.720 | 0.609 | 0.701 | 0.678 |
| | 0.697 | 0.604 | 0.700 | 0.676 |
| | 0.686 | 0.599 | 0.699 | 0.676 |
| | 0.680 | 0.598 | 0.698 | 0.674 |
| | 0.673 | 0.595 | 0.698 | 0.661 |
| | 0.668 | 0.574 | 0.693 | 0.654 |
| | 0.647 | 0.570 | 0.683 | 0.649 |
| | 0.644 | 0.558 | 0.680 | 0.647 |
| | 0.643 | 0.536 | 0.646 | 0.643 |
| | 0.627 | 0.532 | 0.640 | 0.610 |
| AVERAGE ± STD | | | | |
| | 0.791±0.063 | 0.712±0.081 | 0.784±0.060 | 0.767±0.063 |

#### Supplementary Material 5

**Table 1.** Single-cell segmentation tools used in this work

| | **3D-Cell-Annotator (version 1.0)** | **MINS (version 1.3)** | **Pagita (version 2.2)** | **XPIWIT (version 1.0)** | **OpenSegSPIM (version 1.1)** |
|---|---|---|---|---|---|
| DOCUMENTATION | | | | | |
| User guide | ● | ● | ● | ● | ● |
| Website | ● | ● | o | o | ● |
| Video tutorial | ● | o | o | o | ● |
| Freely available tool | ● | ● | ● | ● | ● |
| Open source code | ● | ● | ● | ● | ● |
| Implementation language | C++ | MATLAB/C++ | Java | C++ | MATLAB |
| Test dataset/demo | ● | ● | ● | ● | ● |
| Link to code/executable | www.3d-cell-annotator.org | http://katlab-tools.org | http://imagejdocu.tudor.lu | https: //bitbucket.org/jstegmaier /xpiwit/downloads/ | opensegspim.weebly.com |
| Scientific reference | Tasnadi et al., 2019 | Lou et al., 2014 | Gul-Mohammed et al., 2014 | Bartschat et al., 2015 | Gole et al., 2016 |
| USABILITY | | | | | |
| No programming experience is required | ● | ● | o | o | ● |
| User-friendly GUI | ● | ● | o | o | ● |
| Intuitive visualization settings | ● | ● | o | o | o |
| No commercial license is required | ● | ● | ● | ● | ● |
| Portability on Win/Linux/Mac | Win/Linux | Win | Win/Linux/Mac | Win/Linux/Mac | Win/Mac |
| FUNCTIONALITY | | | | | |

| | | | | | |
|---|---|---|---|---|---|
| Automatic single-cell segmentation | ● | ● | ● | ● | ● |
| Manual correction opportunity | ● | o | ● | o | ● |
| Feature extraction | o | ● | ● | ● | ● |
| No human interaction is required | o | o | o | o | o |
| OUTPUT | | | | | |
| 3D rendering | ● | o | o | ● | o |
| 3D binary mask | ● | ● | ● | ● | ● |
| Features statistics | o | ● | ● | ● | ● |

● available/yes; o not available/no

In this work we compared the segmentations obtained with 3D-Cell-Annotator to other four publicly available tools. Hereafter we give a brief description of these tools.

**1) MINS (version 1.3)**

MINS stands for Modular Interactive Nuclear Segmentation, and it is a MATLAB/C++-based segmentation tool tailored for fluorescent intensity measurements of 2D and 3D image data. Today MINS (Lou *et al.* 2014) is one of the most widely used freely available tools for segmenting single-cells in embryos, but it can easily be used for 3D -oids as well. The MINS pipeline comprises three major cascaded modules: detection, segmentation, and cell position classification. Detection is based on a multiscale blob detection technique, and provides accurate localization of cell nuclei only. Segmentation expands detection output to cover the full nuclear body. The authors chose Seeded Geodesic Image Segmentation (SGIS) as the base algorithm for this stage. Finally, classification is obtained through a clustering-based approach combined with a robust shape-fitting approach and it serves multiple purposes, including the separation of multiple embryos and removal of outliers, as well as the classification of inner and outer cells. However, in practice, MINS requires only a few mouse clicks and a minimum of parameter tuning from the user. The source-code and a Windows-only standalone executable of MINS version 1.3 are distributed at: http://katlab-tools.org.

**2) Pagita (version 2.2)**

Pagita (Gul-Mohammed *et al.* 2014) is an automatic algorithm to segment and simultaneously classify cells/nuclei in 3D/4D images. Segmentation relies on training samples that are interactively provided by the user and in an iterative thresholding process. This algorithm can segment cells/nuclei even when they are touching, and remains effective under temporal and spatial intensity variations. However, it may be quite time-consuming since many threshold values are tested. The algorithm has been implemented as an open-source plug-in for ImageJ and is publicly available for download along with a tutorial and sample data at: http://imagejdocu.tudor.lu. The main idea of the algorithm is to use machine learning to improve the segmentation results. The learning step is an interactive process. The user first selects representative cells/nuclei by clicking on their locations inside the 3D or 4D dataset. The objects at the clicked positions are then segmented through an iterative thresholding procedure using the user-supplied volume estimates as a reference. Next, the user has to validate the proposed segmented nuclei, and 3D descriptors are computed from this set of validated nuclei. The main joint segmentation/classification procedure is subsequently applied to all time points.

**3) XPIWIT (version 1.0)**

XPIWIT (Bartschat *et al*. 2015) stands for XML Pipeline Wizard for ITK. It is an XML-based wrapper application for the Insight Toolkit that combines the performance of a pure C++ implementation with an easy-to-use graphical setup of dynamic image analysis pipelines. The core of XPIWIT is the Threshold of Weighted intensity And seed-Normal Gradient dot product image (TWANG) segmentation (Stegmaier *et al*. 2014). The current version of XPIWIT (*i.e.* version 1.0) incorporates about 70 different filters, and can be constantly extended with new functionality using the provided template files that facilitate the implementation of new modules. To apply a predefined XML processing pipeline to a desired set of images, XPIWIT has to be executed from the command prompt with command line input arguments. Alternatively, a configuration text file can be piped to the executable. This configuration file needs to contain the output path, one or more input paths, the path to the XML file to be processed, and may be customized using further optional parameters as described in the provided documentation. XPIWIT also has an easy-to-use GUI that allows to create and modify XML pipelines based on the filters compiled into the XPIWIT executable. XPIWIT has been successfully applied to segment and visualize terabyte-scale 3D+time light-sheet microscopy images of developing embryos. Its source-code, documentations and standalone versions are available at: https://bitbucket.org/jstegmaier/xpiwit/downloads/.

**4) OpenSegSPIM (version 1.1)**

OpenSegSPIM (Gole *et al*., 2016) is an open-source and user-friendly 3D automatic quantitative analysis tool for confocal/multiphoton/LSFM data. It is a tool developed in MATLAB (The MathWorks, Inc., MA, USA). Its source-code, standalone versions (for Windows and Mac), documentation, user manuals and video tutorials are distributed at: http://opensegspim.weebly.com/. OpenSegSPIM assumes no prior knowledge of image processing or programming and is designed to easily segment nuclei/cells and compute several features without requiring human interaction, except for setting a few initial parameters: an approximate nuclei diameter measurement and intensity adjustment of the image contrast. OpenSegSPIM also includes: (a) a sub-cellular segmentation tool to analyse subcellular compartments, such as the cell membrane; (b) a simple post-processing tool for manually editing the segmentations obtained; and (c) an automatic batch process of time series and numerous datasets. However, the final segmentation of single-cells lacks quality, mainly due to the limits of thresholding in case of blurry objects and an improper separation of touching cells. The currently available version of OpenSegSPIM is version 1.1.

### Supplementary Material 6

**Table 2.** Spheroid dataset: single-cell Jaccard Index (JI) values

| | Annotator 1 vs Annotator 2 | Annotator 1 vs 3D-Cell-Annotator | Annotator 2 vs 3D-Cell-Annotator | Annotator 1 vs MINS | Annotator 2 vs MINS | Annotator 1 vs Pagita | Annotator 2 vs Pagita | Annotator 1 vs XPIWIT | Annotator 2 vs XPIWIT | Annotator 1 vs OpenSegSPIM | Annotator 2 vs OpenSegSPIM |
|---|---|---|---|---|---|---|---|---|---|---|---|
| SINGLE CELL JI | | | | | | | | | | | |
| | 0.887 | 0.889 | 0.900 | 0.856 | 0.722 | 0.840 | 0.834 | 0.745 | 0.625 | 0.735 | 0.696 |
| | 0.830 | 0.859 | 0.872 | 0.763 | 0.700 | 0.756 | 0.611 | 0.743 | 0.616 | 0.730 | 0.653 |
| | 0.819 | 0.837 | 0.855 | 0.763 | 0.667 | 0.747 | 0.577 | 0.718 | 0.609 | 0.703 | 0.584 |
| | 0.818 | 0.836 | 0.847 | 0.761 | 0.620 | 0.677 | 0.543 | 0.706 | 0.591 | 0.700 | 0.564 |
| | 0.789 | 0.822 | 0.837 | 0.733 | 0.610 | 0.659 | 0.474 | 0.704 | 0.588 | 0.693 | 0.554 |
| | 0.788 | 0.813 | 0.813 | 0.726 | 0.606 | 0.639 | 0.457 | 0.692 | 0.575 | 0.671 | 0.550 |
| | 0.784 | 0.813 | 0.806 | 0.724 | 0.606 | 0.533 | 0.435 | 0.690 | 0.572 | 0.660 | 0.532 |
| | 0.781 | 0.811 | 0.801 | 0.713 | 0.598 | 0.528 | 0.423 | 0.689 | 0.554 | 0.646 | 0.525 |
| | 0.781 | 0.811 | 0.799 | 0.709 | 0.583 | 0.464 | 0.365 | 0.689 | 0.552 | 0.640 | 0.518 |
| | 0.774 | 0.796 | 0.798 | 0.707 | 0.576 | 0.458 | 0.363 | 0.685 | 0.536 | 0.620 | 0.508 |
| | 0.774 | 0.782 | 0.797 | 0.688 | 0.574 | 0.442 | 0.360 | 0.676 | 0.533 | 0.620 | 0.505 |
| | 0.770 | 0.781 | 0.797 | 0.686 | 0.564 | 0.441 | 0.343 | 0.669 | 0.526 | 0.612 | 0.502 |
| | 0.760 | 0.779 | 0.794 | 0.685 | 0.549 | 0.441 | 0.317 | 0.666 | 0.524 | 0.608 | 0.500 |
| | 0.745 | 0.778 | 0.792 | 0.683 | 0.548 | 0.430 | 0.315 | 0.661 | 0.521 | 0.606 | 0.500 |
| | 0.741 | 0.776 | 0.788 | 0.678 | 0.540 | 0.427 | 0.314 | 0.658 | 0.521 | 0.605 | 0.496 |
| | 0.737 | 0.774 | 0.786 | 0.670 | 0.530 | 0.396 | 0.310 | 0.651 | 0.517 | 0.605 | 0.484 |
| | 0.729 | 0.766 | 0.784 | 0.666 | 0.527 | 0.394 | 0.305 | 0.644 | 0.517 | 0.602 | 0.484 |
| | 0.724 | 0.765 | 0.781 | 0.666 | 0.525 | 0.392 | 0.291 | 0.640 | 0.509 | 0.595 | 0.483 |

| | | | | | | | | | | |
|---|---|---|---|---|---|---|---|---|---|---|
| 0.724 | 0.760 | 0.781 | 0.657 | 0.524 | 0.376 | 0.289 | 0.630 | 0.505 | 0.580 | 0.468 |
| 0.715 | 0.756 | 0.778 | 0.638 | 0.522 | 0.372 | 0.284 | 0.629 | 0.481 | 0.577 | 0.462 |
| 0.713 | 0.755 | 0.773 | 0.636 | 0.518 | 0.325 | 0.275 | 0.627 | 0.480 | 0.558 | 0.458 |
| 0.704 | 0.750 | 0.764 | 0.631 | 0.516 | 0.323 | 0.273 | 0.617 | 0.476 | 0.557 | 0.458 |
| 0.699 | 0.741 | 0.745 | 0.629 | 0.512 | 0.318 | 0.261 | 0.617 | 0.472 | 0.553 | 0.447 |
| 0.698 | 0.736 | 0.741 | 0.618 | 0.509 | 0.318 | 0.258 | 0.602 | 0.469 | 0.553 | 0.444 |
| 0.682 | 0.734 | 0.738 | 0.613 | 0.488 | 0.314 | 0.255 | 0.592 | 0.453 | 0.553 | 0.434 |
| 0.682 | 0.732 | 0.734 | 0.602 | 0.483 | 0.314 | 0.246 | 0.588 | 0.447 | 0.542 | 0.427 |
| 0.679 | 0.729 | 0.734 | 0.601 | 0.482 | 0.289 | 0.242 | 0.585 | 0.440 | 0.534 | 0.426 |
| 0.676 | 0.723 | 0.734 | 0.597 | 0.478 | 0.289 | 0.232 | 0.585 | 0.439 | 0.514 | 0.424 |
| 0.672 | 0.723 | 0.709 | 0.586 | 0.477 | 0.267 | 0.220 | 0.582 | 0.435 | 0.513 | 0.382 |
| 0.665 | 0.719 | 0.704 | 0.586 | 0.473 | 0.256 | 0.210 | 0.580 | 0.429 | 0.502 | 0.372 |
| 0.660 | 0.718 | 0.698 | 0.581 | 0.453 | 0.233 | 0.208 | 0.575 | 0.415 | 0.501 | 0.367 |
| 0.659 | 0.713 | 0.696 | 0.578 | 0.444 | 0.231 | 0.195 | 0.571 | 0.414 | 0.487 | 0.356 |
| 0.654 | 0.702 | 0.695 | 0.573 | 0.438 | 0.228 | 0.174 | 0.563 | 0.404 | 0.483 | 0.355 |
| 0.652 | 0.694 | 0.690 | 0.571 | 0.427 | 0.190 | 0.174 | 0.561 | 0.400 | 0.472 | 0.350 |
| 0.650 | 0.688 | 0.680 | 0.566 | 0.423 | 0.187 | 0.165 | 0.560 | 0.399 | 0.471 | 0.347 |
| 0.636 | 0.688 | 0.675 | 0.564 | 0.419 | 0.182 | 0.163 | 0.535 | 0.392 | 0.463 | 0.329 |
| 0.635 | 0.670 | 0.605 | 0.549 | 0.405 | 0.168 | 0.156 | 0.518 | 0.390 | 0.459 | 0.325 |
| 0.623 | 0.667 | 0.595 | 0.531 | 0.394 | 0.163 | 0.148 | 0.517 | 0.369 | 0.454 | 0.321 |
| 0.621 | 0.663 | 0.579 | 0.512 | 0.394 | 0.140 | 0.130 | 0.517 | 0.343 | 0.452 | 0.320 |
| 0.619 | 0.644 | 0.560 | 0.497 | 0.376 | 0.135 | 0.113 | 0.517 | 0.341 | 0.447 | 0.316 |

| | | | | | | | | | | | |
|---|---|---|---|---|---|---|---|---|---|---|---|
| | 0.580 | 0.643 | 0.556 | 0.490 | 0.366 | 0.131 | 0.106 | 0.506 | 0.324 | 0.417 | 0.312 |
| | 0.578 | 0.610 | 0.551 | 0.444 | 0.342 | 0.128 | 0.106 | 0.504 | 0.315 | 0.411 | 0.301 |
| | 0.576 | 0.607 | 0.549 | 0.417 | 0.328 | 0.124 | 0.090 | 0.313 | 0.280 | 0.409 | 0.294 |
| | 0.568 | 0.547 | 0.549 | 0.404 | 0.317 | 0.105 | 0.087 | 0.239 | 0.055 | 0.404 | 0.284 |
| | 0.567 | 0.511 | 0.543 | 0.394 | 0.311 | 0.105 | 0.082 | 0.063 | 0.042 | 0.376 | 0.282 |
| | 0.555 | 0.487 | 0.542 | 0.362 | 0.296 | 0.101 | 0.075 | 0.048 | 0.029 | 0.349 | 0.277 |
| | 0.454 | 0.465 | 0.538 | 0.329 | 0.268 | 0.098 | 0.072 | 0.045 | 0.027 | 0.335 | 0.250 |
| | 0.446 | 0.413 | 0.498 | 0.308 | 0.265 | 0.095 | 0.052 | 0.031 | 0.025 | 0.327 | 0.209 |
| | 0.425 | 0.356 | 0.330 | 0.301 | 0.042 | 0.071 | 0.038 | 0.030 | 0.014 | 0.324 | 0.202 |
| | 0.276 | 0.338 | 0.321 | 0.026 | 0.019 | 0.031 | 0.026 | 0.014 | 0.011 | 0.263 | 0.199 |
| | 0.214 | 0.321 | 0.156 | 0.000 | 0.009 | 0.004 | 0.005 | 0.005 | 0.000 | 0.236 | 0.190 |
| | 0.212 | 0.316 | 0.015 | 0.000 | 0.005 | 0.000 | 0.000 | 0.000 | 0.000 | 0.158 | 0.081 |
| AVERAGE ± STD | | | | | | | | | | | |
| | 0.658±0.143 | 0.689±0143 | 0.677±0.175 | 0.563±0.185 | 0.449±0.164 | 0.313±0.203 | 0.251±0.167 | 0.515±0.228 | 0.394±0.189 | 0.517±0.130 | 0.406±0.126 |

#### Supplementary Material 7

**Table 3.** Embryo dataset: single-cell Jaccard Index (JI) values

| | **Annotator 1 vs Annotator 2** | **Annotator 1 vs 3D-Cell-Annotator** | **Annotator 2 vs 3D-Cell-Annotator** |
|---|---|---|---|
| SINGLE CELL JI | | | |
| | 0.852 | 0.881 | 0.771 |
| | 0.721 | 0.794 | 0.746 |
| | 0.771 | 0.881 | 0.762 |
| | 0.809 | 0.889 | 0.827 |
| | 0.836 | 0.893 | 0.856 |
| | 0.789 | 0.888 | 0.853 |
| | 0.845 | 0.874 | 0.665 |
| | 0.878 | 0.896 | 0.768 |
| | 0.831 | 0.869 | 0.871 |
| | 0.766 | 0.891 | 0.778 |
| | 0.789 | 0.871 | 0.830 |
| | 0.858 | 0.878 | 0.810 |
| | 0.875 | 0.909 | 0.826 |
| | 0.684 | 0.832 | 0.827 |
| | 0.802 | 0.761 | 0.841 |
| | 0.784 | 0.908 | 0.805 |
| | 0.820 | 0.849 | 0.802 |
| | 0.796 | 0.862 | 0.852 |

| | | |
|---|---|---|
| 0.804 | 0.843 | 0.825 |
| 0.778 | 0.918 | 0.805 |
| 0.839 | 0.869 | 0.861 |
| 0.659 | 0.846 | 0.812 |
| 0.869 | 0.888 | 0.822 |
| 0.770 | 0.844 | 0.798 |
| 0.815 | 0.797 | 0.727 |
| 0.785 | 0.884 | 0.732 |
| 0.713 | 0.909 | 0.754 |
| 0.727 | 0.866 | 0.821 |
| 0.855 | 0.781 | 0.742 |
| 0.724 | 0.709 | 0.842 |
| 0.803 | 0.768 | 0.837 |
| 0.702 | 0.791 | 0.799 |
| 0.821 | 0.869 | 0.793 |
| 0.827 | 0.853 | 0.688 |
| 0.759 | 0.804 | 0.813 |
| 0.788 | 0.696 | 0.816 |
| 0.690 | 0.752 | 0.747 |
| 0.759 | 0.774 | 0.791 |
| 0.824 | 0.721 | 0.862 |
| 0.834 | 0.625 | 0.795 |

`

| | | | |
|---|---|---|---|
| | 0.842 | 0.775 | 0.762 |
| | 0.682 | 0.671 | 0.740 |
| | 0.760 | 0.647 | 0.725 |
| | 0.765 | 0.666 | 0.770 |
| | 0.767 | 0.677 | 0.880 |
| | 0.586 | 0.584 | 0.737 |
| | 0.623 | 0.627 | 0.855 |
| | 0.596 | 0.702 | 0.759 |
| | 0.706 | 0.718 | 0.871 |
| | 0.797 | 0.697 | 0.880 |
| | 0.809 | 0.767 | 0.868 |
| | 0.857 | 0.783 | 0.882 |
| | 0.770 | 0.785 | 0.841 |
| | 0.790 | 0.800 | 0.794 |
| | 0.844 | 0.720 | 0.724 |
| | 0.791 | 0.878 | 0.681 |
| AVERAGE ± STD | | | |
| | 0.779±0.068 | 0.802±0.088 | 0.799±0.054 |

#### Supplementary Material 8

All the masks and segmentations considered in this work can be freely downloaded at:

*http://www.3d-cell-annotator.org/download.html*
